# Drug-Target-Interaction Prediction with Contrastive and Siamese Transformers

**DOI:** 10.1101/2023.10.26.564262

**Authors:** Daniel Ikechukwu, Arav Kumar

**Affiliations:** Blackbox Research, Manchester, England; Blackbox Research, Palo Alto, California, USA

## Abstract

As machine learning (ML) becomes increasingly integrated into the drug development process, accurately predicting Drug-Target Interactions (DTI) becomes a necessity for pharmaceutical research. This prediction plays a crucial role in various aspects of drug development, including virtual screening, repurposing of drugs, and proactively identifying potential side effects. While Deep Learning has made significant progress in enhancing DTI prediction, challenges related to interpretability and consistent performance persist in the field. This study introduces two innovative methodologies that combine Generative Pretraining and Contrastive Learning to specialize Transformers for biochemical modeling. These systems are designed to best incorporate cross-attention, which enables a nuanced alignment of multi-representation embeddings. Our empirical evaluation will showcase the effectiveness and interpretability of this proposed framework. Through a series of experiments, we provide compelling evidence of its superior predictive accuracy and enhanced interpretability. The primary objective of this research is not only to contribute to the advancement of novel DTI prediction methods but also to promote greater transparency and reliability within the drug discovery pipeline.

## 1 Introduction

In the domain of Drug-Target Interaction (DTI) prediction, the use of Deep Learning has yielded remarkable outcomes. These methods have showcased high predictive accuracy and efficiency relative to previous approaches. [1, 2, 3, 4] We deviate from the typical approach of learning a unified joint embedding representation. Instead, we embrace an alignment-centric framework. This shift in perspective reframes the problem as a matter of “alignment,” with the specific goal of identifying accurate correspondences between segments of a drug’s SMILE sequence and their counterparts in the amino acid sequence of a protein target.

To ensure precise alignment of multiple representations with analytical precision, we employ Cross Attention, a widely acknowledged technique renowned for its effectiveness in multi-representation alignment. [5, 6, 7, 8, 9]. Importantly, the inclusion of the Attention Mechanism within our framework not only serves functional purposes but also augments interpretability [10, 11].

Recent empirical findings underscore the potential of generative pretraining in bolstering downstream model fine-tuning, even under the constraints of limited data[12, 13, 14]. Therefore, our research undertakes the task of cultivating generalizable embeddings via generative pretraining, tapping into the reservoir of versatile and high-quality features accrued during this preparatory phase. These adaptable embeddings, characterized by their inherent malleability, are subsequently repurposed for the formidable task of DTI prediction.

The historical non-machine learning based methods have been lacking in all measures of performance relative to what this new technology is capable of. Below, we derive in greater detail the shortcomings of each individual method, but it is noted that on average, the biologically-inspired methods prove less performative.

## 2 Background

The identification of drug-target interactions (DTIs) represents a pivotal and enduring challenge in the ever-evolving field of drug discovery and development.

Over the years, the complexity surrounding DTI prediction has spurred the development of numerous computational methods and approaches. These approaches aim to unravel the intricate web of molecular interactions between drugs and their target proteins, paving the way for informed drug design and optimization.

The existing approaches in DTI prediction can be classified into four major categories: Ligand-based approaches, Target-based approaches, Network-based approaches and Machine Learning-based approaches.

### 2.1 Ligand-based Approaches

Ligand-based methods focus on the molecular properties of drugs and their chemical similarity to known ligands. [15].

Ligand-based methods have indeed demonstrated remarkable utility in predicting drug-target interactions (DTIs) [16, 17, 18]; however, they are not without limitations. One of the most prominent shortcomings is their limited ability to generalize to compounds that differ significantly in structural characteristics from known ligands. [19, 20, 21].

When novel compounds possess structural features that deviate substantially from the training set of known ligands, ligand-based methods struggle to make accurate predictions. These methods rely heavily on the similarity principle, assuming that compounds with similar chemical structures exhibit similar bioactivity profiles. Consequently, the predictive power of ligand-based models diminishes when confronted with structurally dissimilar compounds, rendering them less effective for exploring chemical space beyond the scope of the training data.

### 2.2 Target-based Approaches

Target-based methods in drug-target interaction (DTI) prediction primarily center around the biological characteristics and properties of the drug target itself [22, 23]. These approaches have gained considerable attention in recent years, offering a different perspective compared to ligand-based methods.

Target-based methods leverage knowledge about the specific target proteins or biomolecules involved in drug interactions. They often rely on various biological data sources, such as protein sequences, structures, and functional annotations, to make predictions about potential drug-target interactions [24].

One of the key advantages of target-based approaches is their potential to overcome the limitations of ligand-based methods. Unlike ligand-based methods that struggle with structurally dissimilar compounds, target-based approaches can provide insights into how a particular drug interacts with a specific biological target [25]. This can be particularly valuable when dealing with novel compounds or when there is limited structural similarity to known ligands [26].

However, it’s essential to acknowledge that target-based approaches also come with their set of challenges. They often require extensive biological data, including high-quality structural information about the target proteins [27, 28]. Obtaining such data can be time-consuming and costly. Additionally, the reliability of target-based predictions can be affected by the quality of the available biological data and the accuracy of the target annotation [29, 22].

### 2.3 Network-based Approaches

Network-based approaches represent a promising avenue for predicting Drug-Target Interactions (DTIs) by leveraging the inherent connectivity and interaction patterns within biological systems [30, 31]. These methods capitalize on the idea that proteins and drugs can be viewed as nodes in a complex network, with edges representing interactions or relationships. By analyzing these networks, network-based approaches offer a unique perspective on DTIs, often revealing hidden associations that are challenging to uncover using traditional methods.

One of the key advantages of network-based approaches is their ability to capture the contextual information surrounding a drug or a target [32]. Instead of solely relying on the structural features of molecules or ligands, these methods consider the broader biological context. For instance, they can incorporate protein-protein interaction networks, gene expression data, and pathway information. This holistic view enables network-based models to make predictions that are robust to structural variations in compounds and to better generalize to novel compounds or targets, which can be particularly valuable when dealing with the vast and diverse chemical space of potential drugs.

However, network-based approaches also face their own set of challenges. The quality and completeness of the biological networks used as a basis for prediction can significantly impact the accuracy of DTI predictions [33]. Additionally, interpreting the results of these models can be complex, as they often produce large networks of potential interactions leaving biologists to verify their success. [33].

### 2.4 Machine Learning Approaches

There has been a surge in machine learning based approaches which has demonstrated ML as a powerful tool to complement traditional methods. Target-based approaches focus on the characteristics of protein targets and how they interact with potential drug compounds. These methods have gained significant attention due to their ability to leverage large-scale biological and chemical data, allowing for more comprehensive predictions.

One notable class of these techniques that has shown promise in DTI prediction is contrastive learning. Contrastive learning techniques, often rooted in deep learning, have been employed to capture intricate relationships between drug compounds and protein targets. These approaches involve training models to distinguish between positive and negative pairs of interactions. By learning to differentiate true interactions from non-interactions, these models can effectively represent the underlying patterns in DTIs. Studies have demonstrated the efficacy of contrastive learning in improving the accuracy of DTI predictions [34, 35, 36].

Transformers, originally designed for natural language processing tasks, have found their way into the field of DTI prediction. These models excel at capturing long-range dependencies and complex relationships within sequences of data, making them well-suited for tasks involving biomolecular interactions. Researchers have adapted transformer architectures to encode both drug and target representations, enabling the prediction of DTIs. The ability of transformers to process information in parallel has proven valuable in capturing the nuances of drug-protein interactions [37, 38].

Pretrained models, often fine-tuned on DTI-specific data, have also gained prominence [39]. These models, pretrained on vast amounts of chemical and biological information, can effectively transfer their knowledge to DTI prediction tasks. By leveraging pretrained representations, researchers can significantly enhance the generalization capabilities of DTI models. This transfer learning paradigm has shown promise in improving the accuracy of predictions, especially for rare or poorly characterized drug-target interactions [40, 41, 41].

Deep neural networks remain at the forefront of DTI prediction. These models can capture complex, non-linear relationships between drug compounds and protein targets. Researchers continue to innovate by developing novel neural network architectures tailored to DTI prediction. Deep learning techniques, combined with large-scale biological datasets, have the potential to revolutionize our understanding of drug-target interactions and expedite the drug discovery process [42, 43, 43, 44].

## 3 Methods

### 3.1 Problem Definition

We pose the Drug-Target Interaction problem as a binary classification problem; determining whether or not a drug and protein-target will interact. Following [45], we represent a drug by it’s Simplified Molecular Input Line Entry System (SMILES) string 𝒮 = {*T*_1_,*T*_2_, …, *T*_*t*_} and a protein target by it’s amino-acids sequence 𝒜 = {*P*_1_, *P*_2_,…, *P*_*k*_}. P represents the 23 amino-acids tokens and T represents the list of ASCII characters used in the SMILES representation.

We use two functions, *h*_*drug*_ and *h*_*amino*_ to encode a given SMILE sequence 𝒮 and amino-acids sequence 𝒜 and respectively into latent representations **L**_s_ ∈ ℝ^*n*^ and **L**_a_ ∈ ℝ^*m*^ defined below.

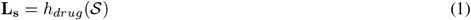

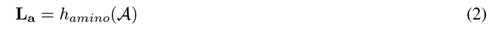

With this, we can determining whether or not a drug-target pair will interact by discovering what aspects of a given drug correlate to it’s ability to interact with the protein-target.

The model, *f* created to represent these latent representations produces the probability of a succesful interaction given the drug-target pair and is represented as follows.

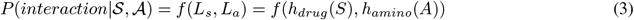

### 3.2 Generative Pretraining For Embedding Generation

Generative pretraining is a well-established technique in natural language processing (NLP) that involves training a language model on a substantial corpus of text data in an unsupervised manner. This pretraining phase equips the model with a foundation to learn intricate representations of natural language. These acquired representations can subsequently be adapted for specific downstream tasks, including sentiment analysis, named entity recognition, and machine translation. [46, 47, 48, 49] The success of generative pretraining in natural language processing can be attributed to its ability to learn rich and meaningful representations of language that can be used for a variety of downstream tasks.

In our approach, we train a simple sequence model in an unsupervised manner. We train it to maximise the log-likelihood of predicting an amino acid given the previous amino acids in the sequence for protein targets. We also train it to predict the next SMILE character given the previous SMILE characters in the case of drugs.

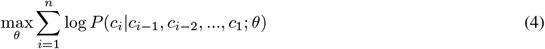

Following the generative pretraining phase, we extract and freeze the learned embeddings as pretrained embeddings within our classification models. The generative pretraining step is visually represented in Figure **??**

### 3.3 Utilizing Contrastive Optimization to Model Transformers as Siamese Networks

Siamese networks represent a unique category of neural networks that undergo training either through contrastive methods or in a purely supervised fashion. These networks are designed to assess the similarity between two input entities. Siamese networks have found extensive application across various domains, ranging from their utilization in tasks such as identifying signature forgery [50] or more broadly, pair anomaly detection [51, 52] to their role in enhancing the efficiency of one-shot classification systems [53, 54].

In the context of neural network architectures, it is noteworthy that transformer models can be employed as siamese networks. In this configuration, the encoder and decoder components are tasked with processing distinct inputs, and the determination of similarity is achieved through cross-attention mechanisms.

This approach eliminates the need for a shared representation space, focusing on the precise alignment of diverse representations, such as those for SMILE sequences and amino acid sequences.

Our contribution involves a unique adaptation of the encoder-decoder transformer architecture, introducing separate inputs to the encoder and decoder components, thereby refining the conventional transformer framework. This evolution enhances the versatility of the transformer model, offering new possibilities in neural network design.

Although Siamese networks utilise shared parameters in learning to differentiate between inputs, we repurpose an Encoder-Decoder Transformer with the same purpose as a Siamese network. We do this by changing the objective from learning to quantify similarity from the difference in representation to determining the aspects of the representations that are in mutual agreeance.

In our approach, we introduce a subtly distinct neural network architecture, as illustrated in Figure 3. This network is made to acheive a dual objective: It is trained to reduce the dissimilarity between latent representations of drugs and targets that possess a well-established history of interaction while at the same time amplifying the contrast in latent representations for those pairs where interactions are unlikely.

**Figure 1.**
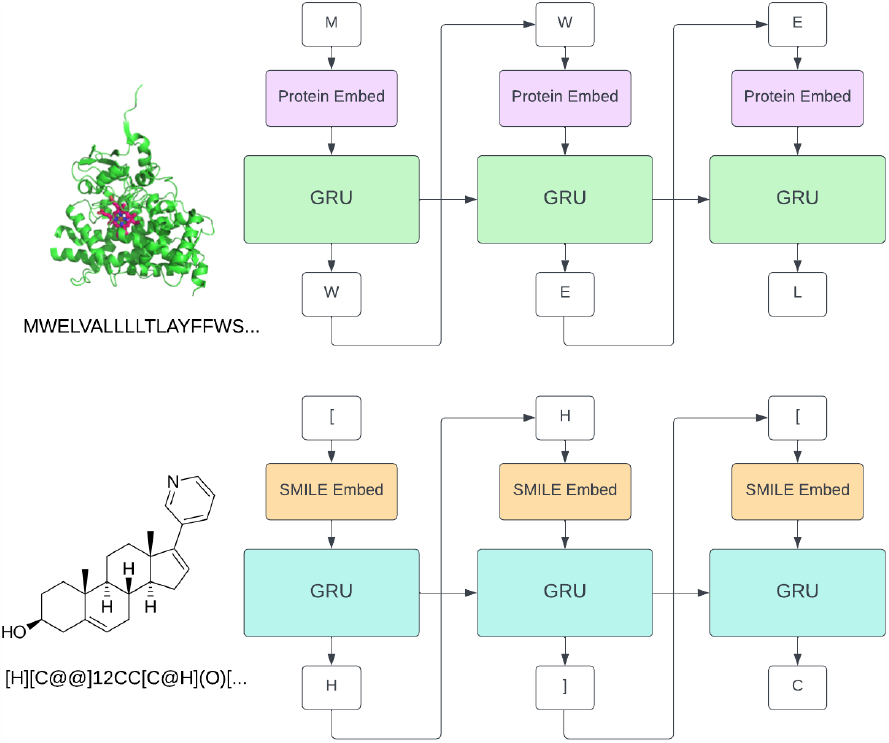
Encoder-Decoder Transformer architecture.

**Figure 2.**
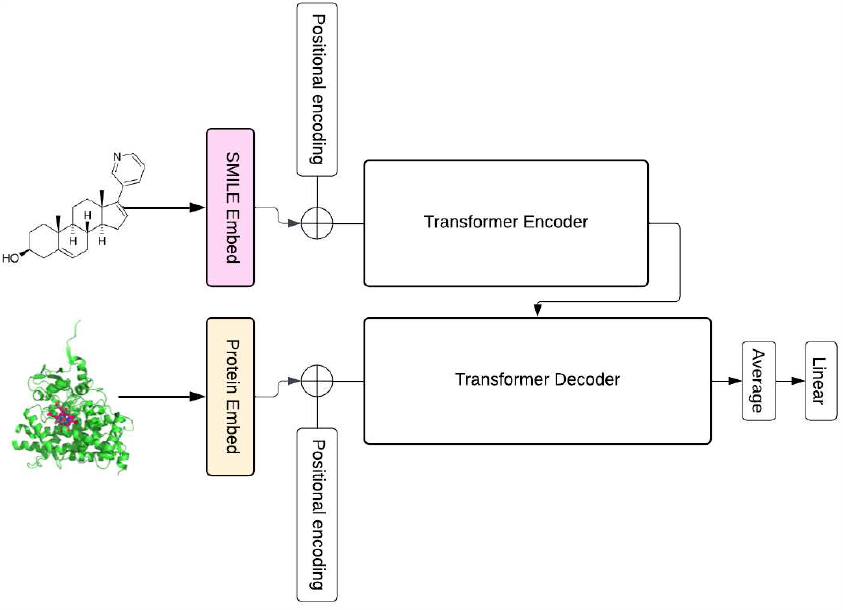
Encoder-Decoder Transformer architecture.

**Figure 3.**
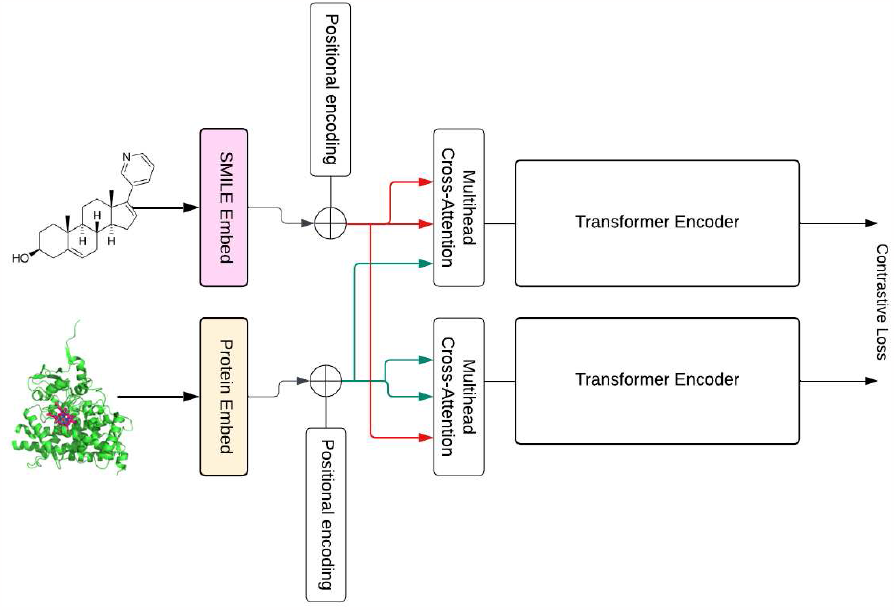
Contrastively optimised Cross-Attention Encoder

The network is optimised by minimising the following contrastive loss function:

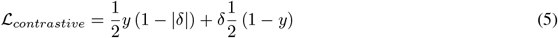

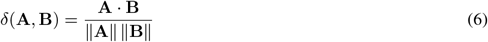

## 4 Results

### 4.1 Setup

We make use of 4 popular datasets for DTI prediction namely BIOSNAP [55], DAVIS [56], BindingDB [57] and KIBA [58].

As the BIOSNAP dataset only contains positive pairs, we produce negative examples by sampling interactions that do not appear in the positive pairs thereby giving us a balanced dataset with equal proportions of positive and negative interactions.

DAVIS, KIBA and BindingDB contained large class imbalances which we mitigate by oversampling the underrepresented class and undersampling the overrepresented class to ensure overall class balance.

The distribution of the datasets in the combined datasets is shown in Table 1

**Table 1:**
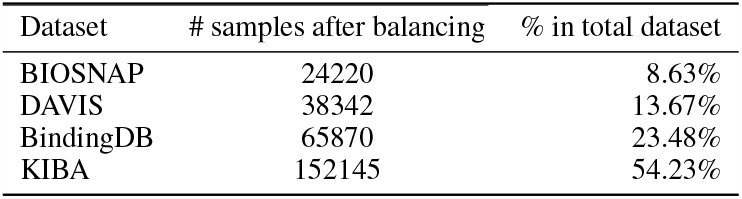
Distribution of datasets.

**Table 2:** Experimentational results.

| Model | Dataset | AUC-ROC |
| --- | --- | --- |
| MolTrans | BIOSNAP + DAVIS + BindingDB + KIBA | 0.905 |
| DNN | BIOSNAP + DAVIS + BindingDB + KIBA | 0.873 |
| DeepDTI | BIOSNAP + DAVIS + BindingDB + KIBA | 0.860 |
| DeepDTA | BIOSNAP + DAVIS + BindingDB + KIBA | 0.890 |
| Ours (Transformer) | BIOSNAP + DAVIS + BindingDB + KIBA | 0.871 |
| Ours (Contrastive Transformer In Progress) | BIOSNAP + DAVIS + BindingDB + KIBA | - |

### 4.2 Implementation details

For our pretrained embeddings sequence models, we train two separate yet closely related Gated Recurrent Units (GRUs). [59] These GRUs are configured with a hidden size *H* = 128, an embedding size *E* = 64, a single layer *N* = 1, and a batch size *B* = 128. The embedding size remains constant across all models as we leverage frozen pretraining from the GRUs.

In the case of our encoder-decoder transformer and contrastive model implementations, we opt for a larger hidden size of *H* = 256 to accommodate the complexity of the tasks. Both models feature two encoder and decoder layers *N* = 2 and benefit from enhanced attention mechanisms with four attention heads *H*_*a*_ = 4. The batch size during training is set at *B* = 8 to balance computational efficiency and model performance.

We maintain a consistent embedding size of *E* = 64 across all our models. This choice is made because we effectively freeze and leverage the embeddings obtained from the pretrained GRUs. This uniformity ensures that our models benefit from the same foundational representation of the input data.

To facilitate model training, we employ the Adam optimizer \cite{kingma2014adam}. Our learning rate *α* = 0.0003 and *β*_1_ & *β*_2_ set to 0.9 and 0.999 respectively.

We utilise dropout [60] to reduce overfitting of the transformer models. We use a dropout value of 0.2 in our encoder-decoder model and a value of 0.1 for our contrastive model. No dropout is used in the training of the GRU models.

### 4.3 Baselines

We test our proposed models against 4 other models namely: MolTrans [45], DNN, DeepDTI [61] and DeepDTA [62].

We report the Area Under the Curve of the “Receiver Operating Characteristic curve (AUC-ROC) of the respective models on the heldout test set.

## 5 Discussion

In this section, we discuss the main findings and implications of our study on Drug-Target Interaction (DTI) prediction using contrastive learning and generative pretraining. Our approach has demonstrated promising results, and we delve into the implications of our findings:

Our experimental results have shown that our novel methodology, which combines generative pretraining with contrastive learning and leverages cross-attention, achieves results on par with other state-of-the-art models and even surpassing some. This indicates that our approach effectively captures the intricate relationships between drugs and protein targets, leading to improved DTI predictions.

The incorporation of cross-attention in our framework not only improves predictive accuracy but also enhances interpretability. This is a significant advancement in the field of DTI prediction, as understanding the basis for predictions is crucial for researchers and practitioners in drug discovery. The attention mechanisms in our model allow us to identify the specific features and interactions that contribute to the prediction, making it more transparent and informative.

One of the persistent challenges in DTI prediction is the limited availability of labeled data. Our use of generative pretraining addresses this issue by providing robust representations even with limited labeled samples. This is especially valuable in scenarios where acquiring large amounts of labeled data is impractical or expensive.

Additionally, we note that this method outperforms past knowledge-based methods of bioinformatics as they don’t rely on incomplete existing theories of biology. This benefit is also paired with an equivalent downside of the limited explainability of these deep learning methods. The limited explainability prevents us from efficiently deriving a true understanding of which elements of a drug interact with their targets.

We have introduced an alignment-focused approach to DTI prediction, diverging from the conventional pursuit of a unified embedding space. By treating DTI prediction as an “alignment” challenge, we highlight the importance of discerning precise correspondences between molecular representations. This approach can be particularly beneficial when dealing with complex drug-target interactions that may involve correlated features within the embeddings.

## 6 Conclusion and Future Work

In conclusion, our study presents a novel framework for DTI prediction that combines generative pretraining, contrastive learning, and cross-attention to achieve enhanced predictive accuracy and interpretability. We have shown that our approach performs competitively with other state-of-the-art models on multiple datasets.

While our current study has yielded promising results, there are several exciting directions for future research that can further advance the field of DTI prediction:

- **Multi-Modal Data Integration:** While our approach leverages molecular representations, future research could explore the integration of additional data modalities such as gene expression profiles, protein-protein interaction networks, and structural biology data. This multi-modal approach may lead to more comprehensive and accurate predictions.
- **Enhanced Interpretability and Transparency:** Enhancing the interpretability of DTI prediction models remains a critical area of research. Future work could focus on developing techniques to provide detailed explanations for model predictions, enabling researchers to gain insights into the biological mechanisms driving interactions.
- **Optimal representation of chemical compound and proteins:** One of the critical aspects of applying sequence models in this line of work is determining the best way to represent chemical compounds and proteins. The choice of how we represent these molecular entities significantly impacts the performance and quality of embeddings produced in the pretraining stage. In future research, optimizing and improving the SMILE representation and other relevant methods for machine learning should be a central focus to further advance the field.

## References

[1] Gert-Jan Bekker, Mitsugu Araki, Kanji Oshima, Yasushi Okuno, and Narutoshi Kamiya. Accurate binding configuration prediction of a g-protein-coupled receptor to its antagonist using multicanonical molecular dynamics-based dynamic docking. Journal of Chemical Information and Modeling, 61(10):5161–5171, 2021.

[2] Annabelle G Vincent and Josh T Beckham. High-performance computational molecular docking for potential inhibitors of an essential enzyme of burkholderia pseudomallei. The FASEB Journal, 36, 2022.

[3] Mohamed Mfa, Sayed Am, Abdelmohsen Ur, N. A., Khashaba Py, and Hayallah Am. Histone deacetylase inhibitors as potential covid-19 virus rna-dependent rna polymerase inhibitors: A molecular docking and dynamics study. Austin - Critical Care Journal, 2021.

[4] Surabhi Jain, Smriti Sharma, Dhrubo Jyoti Sen Surabhi Jain, Smriti Sharma, and Dhrubo Jyoti Sen. Virtual screening, docking, admet and molecular dynamics: A study to find novel inhibitors of mycobacterium tuberculosis targeting qcrb. Jordan Journal of Chemistry (JJC), 16(3):131–146, 2021.

[5] Ashish Vaswani, Noam Shazeer, Niki Parmar, Jakob Uszkoreit, Llion Jones, Aidan N Gomez, Łukasz Kaiser, and Illia Polosukhin. Attention is all you need. Advances in neural information processing systems, 30, 2017.

[6] Binyang Song, Scarlett Miller, and Faez Ahmed. Attention-enhanced multimodal learning for conceptual design evaluations. Journal of Mechanical Design, 145(4):041410, 2023.

[7] Gan Cai, Yu Zhu, Yue Wu, Xiaoben Jiang, Jiongyao Ye, and Dawei Yang. A multimodal transformer to fuse images and metadata for skin disease classification. The Visual Computer, pages 1–13, 2022.

[8] Tadas Baltrušaitis, Chaitanya Ahuja, and Louis-Philippe Morency. Multimodal machine learning: A survey and taxonomy. IEEE transactions on pattern analysis and machine intelligence, 41(2):423–443, 2018.

[9] S Lu, M Liu, L Yin, Z Yin, X Liu, W Zheng, and X Kong. The multi-modal fusion in visual question answering: a review of attention mechanisms. peerj comput sci 9: e1400, 2023.

[10] Shikhar Vashishth, Shyam Upadhyay, Gaurav Singh Tomar, and Manaal Faruqui. Attention interpretability across nlp tasks. arXiv preprint arXiv:1909.11218, 2019.

[11] Lakshmi Narayan Pandey, Rahul Vashisht, and Harish G Ramaswamy. On the interpretability of attention networks. In Asian Conference on Machine Learning, pages 832–847. PMLR, 2023.

[12] Matthew BA McDermott, Brendan Yap, Peter Szolovits, and Marinka Zitnik. Structure-inducing pre-training. Nature Machine Intelligence, pages 1–10, 2023.

[13] Ananya Kumar, Aditi Raghunathan, Robbie Jones, Tengyu Ma, and Percy Liang. Fine-tuning can distort pretrained features and underperform out-of-distribution. arXiv preprint arXiv:2202.10054, 2022.

[14] Ning Ding, Yujia Qin, Guang Yang, Fuchao Wei, Zonghan Yang, Yusheng Su, Shengding Hu, Yulin Chen, Chi-Min Chan, Weize Chen, et al. Parameter-efficient fine-tuning of large-scale pre-trained language models. Nature Machine Intelligence, 5(3):220–235, 2023.

[15] Adel Hamza, Ning-Ning Wei, and Chang-Guo Zhan. Ligand-based virtual screening approach using a new scoring function. Journal of chemical information and modeling, 52(4):963–974, 2012.

[16] Yixian Huang, Hsi-Yuan Huang, Yigang Chen, Yang-Chi-Dung Lin, Lantian Yao, Tianxiu Lin, Junlin Leng, Yuan Chang, Yuntian Zhang, Zihao Zhu, et al. A robust drug–target interaction prediction framework with capsule network and transfer learning. International Journal of Molecular Sciences, 24(18):14061, 2023.

[17] Wail Ba-Alawi, Othman Soufan, Magbubah Essack, Panos Kalnis, and Vladimir B Bajic. Daspfind: new efficient method to predict drug–target interactions. Journal of cheminformatics, 8:1–9, 2016.

[18] Murat Can Cobanoglu, Chang Liu, Feizhuo Hu, Zoltán N Oltvai, and Ivet Bahar. Predicting drug–target interactions using probabilistic matrix factorization. Journal of chemical information and modeling, 53(12):3399–3409, 2013.

[19] Javier Vázquez, Manel López, Enric Gibert, Enric Herrero, and F Javier Luque. Merging ligand-based and structure-based methods in drug discovery: An overview of combined virtual screening approaches. Molecules, 25(20):4723, 2020.

[20] Sam Z Grinter and Xiaoqin Zou. Challenges, applications, and recent advances of protein-ligand docking in structure-based drug design. Molecules, 19(7):10150–10176, 2014.

[21] J Shim and AD Mackerell Jr. Computational ligand-based rational design: role of conformational sampling and force fields in model development. medchemcomm 2: 356–370, 2011.

[22] Qing Ye, Chang-Yu Hsieh, Ziyi Yang, Yu Kang, Jiming Chen, Dongsheng Cao, Shibo He, and Tingjun Hou. A unified drug–target interaction prediction framework based on knowledge graph and recommendation system. Nature communications, 12(1):6775, 2021.

[23] Xuting Zhang, Fengxu Wu, Nan Yang, Xiaohui Zhan, Jianbo Liao, Shangkang Mai, and Zunnan Huang. In silico methods for identification of potential therapeutic targets. Interdisciplinary Sciences: Computational Life Sciences, pages 1–26, 2022.

[24] Zengrui Wu, Weihua Li, Guixia Liu, and Yun Tang. Network-based methods for prediction of drug-target interactions. Frontiers in pharmacology, 9:1134, 2018.

[25] Deborah Giordano, Carmen Biancaniello, Maria Antonia Argenio, and Angelo Facchiano. Drug design by pharmacophore and virtual screening approach. Pharmaceuticals, 15(5):646, 2022.

[26] Prasannavenkatesh Durai, Young-Joon Ko, Cheol-Ho Pan, and Keunwan Park. Evolutionary chemical binding similarity approach integrated with 3d-qsar method for effective virtual screening. BMC bioinformatics, 21:1–18, 2020.

[27] Christoph H Emmerich, Lorena Martinez Gamboa, Martine CJ Hofmann, Marc Bonin-Andresen, Olga Arbach, Pascal Schendel, Björn Gerlach, Katja Hempel, Anton Bespalov, Ulrich Dirnagl, et al. Improving target assessment in biomedical research: the got-it recommendations. Nature reviews Drug discovery, 20(1):64–81, 2021.

[28] John G Moffat, Fabien Vincent, Jonathan A Lee, Jörg Eder, and Marco Prunotto. Opportunities and challenges in phenotypic drug discovery: an industry perspective. Nature reviews Drug discovery, 16(8):531–543, 2017.

[29] Yuanyuan Zhang, Mengjie Wu, Shudong Wang, and Wei Chen. Efmsdti: Drug-target interaction prediction based on an efficient fusion of multi-source data. Frontiers in Pharmacology, 13:1009996, 2022.

[30] Shanglin Gao, Zhixing Liu, and Ying Li. Networks and algorithms in heterogeneous network-based methods for drug-target interaction prediction: A survey and comparison. Proceedings of the 1st International Conference on Health Big Data and Intelligent Healthcare, 2022.

[31] Zhan-Heng Chen, Zhu-Hong You, Zhen-Hao Guo, Hai-Cheng Yi, Gong-Xu Luo, and Yan-Bin Wang. Prediction of drug–target interactions from multi-molecular network based on deep walk embedding model. Frontiers in Bioengineering and Biotechnology, 8:338, 2020.

[32] JM Harrold, M Ramanathan, and DE Mager. Network-based approaches in drug discovery and early development. Clinical Pharmacology & Therapeutics, 94(6):651–658, 2013.

[33] Xiangxiang Zeng, Siyi Zhu, Yuan Hou, Pengyue Zhang, Lang Li, Jing Li, L Frank Huang, Stephen J Lewis, Ruth Nussinov, and Feixiong Cheng. Network-based prediction of drug–target interactions using an arbitrary-order proximity embedded deep forest. Bioinformatics, 36(9):2805–2812, 2020.

[34] Rohit Singh, Samuel Sledzieski, Bryan Bryson, Lenore Cowen, and Bonnie Berger. Contrastive learning in protein language space predicts interactions between drugs and protein targets. Proceedings of the National Academy of Sciences, 120(24):e2220778120, 2023.

[35] Kainan Yao, Xiaowen Wang, Wannian Li, Hongming Zhu, Yizhi Jiang, Yulong Li, Tongxuan Tian, Zhaoyi Yang, Qi Liu, and Qin Liu. Semi-supervised heterogeneous graph contrastive learning for drug–target interaction prediction. Computers in Biology and Medicine, 163:107199, 2023.

[36] Ziduo Yang, Weihe Zhong, Lu Zhao, and Calvin Yu-Chian Chen. Ml-dti: mutual learning mechanism for interpretable drug–target interaction prediction. The Journal of Physical Chemistry Letters, 12(17):4247–4261, 2021.

[37] Sabeen Ahmed, Ian E Nielsen, Aakash Tripathi, Shamoon Siddiqui, Ravi P Ramachandran, and Ghulam Rasool. Transformers in time-series analysis: A tutorial. Circuits, Systems, and Signal Processing, pages 1–34, 2023.

[38] Xiangfeng Yan and Yong Liu. Graph–sequence attention and transformer for predicting drug–target affinity. RSC advances, 12(45):29525–29534, 2022.

[39] Hisham Abdel-Aty and Ian R Gould. Large-scale distributed training of transformers for chemical fingerprinting. Journal of Chemical Information and Modeling, 62(20):4852–4862, 2022.

[40] Jie Zheng, Xuan Xiao, and Wang-Ren Qiu. Dti-bert: identifying drug-target interactions in cellular networking based on bert and deep learning method. Frontiers in Genetics, 13:859188, 2022.

[41] Derwin Suhartono, Muhammad Rizki Nur Majiid, Alif Tri Handoyo, Pandu Wicaksono, and Henry Lucky. Towards a more general drug target interaction prediction model using transfer learning. Procedia Computer Science, 216:370–376, 2023.

[42] Farshid Rayhan, Sajid Ahmed, Zaynab Mousavian, Dewan Md Farid, and Swakkhar Shatabda. Frnet-dti: Deep convolutional neural network for drug-target interaction prediction. Heliyon, 6(3), 2020.

[43] Lu Wang, Yifeng Zhou, and Qu Chen. Ammvf-dti: A novel model predicting drug–target interactions based on attention mechanism and multi-view fusion. International Journal of Molecular Sciences, 24(18):14142, 2023.

[44] Ingoo Lee, Jongsoo Keum, and Hojung Nam. Deepconv-dti: Prediction of drug-target interactions via deep learning with convolution on protein sequences. PLoS computational biology, 15(6):e1007129, 2019.

[45] Kexin Huang, Cao Xiao, Lucas M Glass, and Jimeng Sun. Moltrans: molecular interaction transformer for drug–target interaction prediction. Bioinformatics, 37(6):830–836, 2021.

[46] Alec Radford, Karthik Narasimhan, Tim Salimans, Ilya Sutskever, et al. Improving language understanding by generative pre-training. 2018.

[47] Junyi Li, Tianyi Tang, Wayne Xin Zhao, Jian-Yun Nie, and Ji-Rong Wen. Pretrained language models for text generation: A survey. arXiv preprint arXiv:2201.05273, 2022.

[48] M Ramprasath, K Dhanasekaran, T Karthick, R Velumani, and P Sudhakaran. An extensive study on pretrained models for natural language processing based on transformers. In 2022 International Conference on Electronics and Renewable Systems (ICEARS), pages 382–389. IEEE, 2022.

[49] Thorben Schomacker and Marina Tropmann-Frick. Language representation models: An overview. Entropy, 23(11):1422, 2021.

[50] Ojaswini Chhabra and Souradip Chakraborty. Siamese triple ranking convolution network in signature forgery detection. In Proceedings of the Alliance International Conference on Artificial Intelligence and Machine Learning (AICAAM), 2019.

[51] Shayan Hashemi and Mika Mäntylä. Detecting anomalies in software execution logs with siamese network. arXiv preprint arXiv:2102.01452, 2021.

[52] Niamh Belton, Misgina Tsighe Hagos, Aonghus Lawlor, and Kathleen M Curran. Fewsome: One-class few shot anomaly detection with siamese networks. In Proceedings of the IEEE/CVF Conference on Computer Vision and Pattern Recognition, pages 2977–2986, 2023.

[53] Thomas Müller, Guillermo Pérez-Torró, and Marc Franco-Salvador. Few-shot learning with siamese networks and label tuning. arXiv preprint arXiv:2203.14655, 2022.

[54] Bin Wang and Dian Wang. Plant leaves classification: A few-shot learning method based on siamese network. Ieee Access, 7:151754–151763, 2019.

[55] Kexin Huang, Cao Xiao, Trong Hoang, Lucas Glass, and Jimeng Sun. Caster: Predicting drug interactions with chemical substructure representation. In Proceedings of the AAAI conference on artificial intelligence, volume 34, pages 702–709, 2020.

[56] Mindy I Davis, Jeremy P Hunt, Sanna Herrgard, Pietro Ciceri, Lisa M Wodicka, Gabriel Pallares, Michael Hocker, Daniel K Treiber, and Patrick P Zarrinkar. Comprehensive analysis of kinase inhibitor selectivity. Nature biotechnology, 29(11):1046–1051, 2011.

[57] Tiqing Liu, Yuhmei Lin, Xin Wen, Robert N Jorissen, and Michael K Gilson. Bindingdb: a web-accessible database of experimentally determined protein–ligand binding affinities. Nucleic acids research, 35(suppl_1):D198–D201, 2007.

[58] Jing Tang, Agnieszka Szwajda, Sushil Shakyawar, Tao Xu, Petteri Hintsanen, Krister Wennerberg, and Tero Aittokallio. Making sense of large-scale kinase inhibitor bioactivity data sets: a comparative and integrative analysis. Journal of Chemical Information and Modeling, 54(3):735–743, 2014.

[59] Junyoung Chung, Caglar Gulcehre, KyungHyun Cho, and Yoshua Bengio. Empirical evaluation of gated recurrent neural networks on sequence modeling. arXiv preprint arXiv:1412.3555, 2014.

[60] Nitish Srivastava, Geoffrey Hinton, Alex Krizhevsky, Ilya Sutskever, and Ruslan Salakhutdinov. Dropout: a simple way to prevent neural networks from overfitting. The journal of machine learning research, 15(1):1929–1958, 2014.

[61] Ming Wen, Zhimin Zhang, Shaoyu Niu, Haozhi Sha, Ruihan Yang, Yonghuan Yun, and Hongmei Lu. Deep-learning-based drug–target interaction prediction. Journal of proteome research, 16(4):1401–1409, 2017.

[62] Hakime Öztürk, Arzucan Özgür, and Elif Ozkirimli. Deepdta: deep drug–target binding affinity prediction. Bioinformatics, 34(17):i821–i829, 2018.

